# CROCHET: a versatile pipeline for automated analysis and visual atlas creation from single-cell spatialomic data

**DOI:** 10.64898/2026.03.13.711472

**Authors:** Behnaz Bozorgui, Guillaume Thibault, Chunyu Yuan, Zeynep Dereli, Huamin Wang, Michael J. Overman, John N. Weinstein, Anil Korkut

## Abstract

Spatial biology technologies offer a unique opportunity to link tissue composition with function. However, analytical methods for quantifying and interpreting highly complex spatial data remain limited. We present CROCHET (ChaRacterization Of Cellular HEterogeneity in Tissues), an end-to-end analysis pipeline for construction of spatially resolved cell atlases from raw data covering millions of cells across large sample cohorts. Its modular architecture supports the integration of diverse data modalities and novel analytical methods for image processing and segmentation, spatialomics quantification and downstream analyses. With comprehensive, open-source, user-friendly, interactive, and visual analysis modules, CROCHET aims to democratize spatial omics for a broad community of users.

## Main

Spatial omics technologies hold significant promise for elucidating cellular functions and interactions that mediate key biological processes including cancer progression, immune responses, and cardiovascular diseases^1–3^. These technologies can generate spatial data with unprecedented levels of detail on tissue characteristics. To fully leverage the spatial omics data and extract biologically meaningful insights, there is a major need for robust analytical methods that can perform rigorous quality control, accurate molecular quantitation, and comprehensive downstream analyses.

To address this major challenge, we developed CROCHET, an end-to-end spatial omics analysis pipeline that automates the multi-step transformation from raw multiplexed tissue images to a spatially resolved single-cell atlas. Although optimized for analysis of highly multiplexed imaging-based proteomics data, the pipeline is also applicable to analysis of single-cell spatial transcriptomics data from imaging-based platforms (e.g., 10X Xenium)^4^. In addition to enabling end-to-end analysis, CROCHET introduces a suite of novel computational methods. These include: (i) detection and removal of false positives and residual signals in multiplexed cellular imaging data; (ii) a spatial enrichment score that integrates expression-weighted proximity metrics to capture complex spatial gradients; (iii) multiscale neighborhood detection based on spatial enrichment scores; (iv) distance-aware ligand–receptor modeling that quantifies cell–cell and protein-level interactions beyond binary adjacency or discrete neighborhood definitions, and resulting *immunoprints* for spatial mapping of immunotherapy targets that are recurrent across patient cohorts; (v) an advanced visualization toolkit; and (vii) capabilities for 3D tissue reconstruction. The supplementary table 1 provides a detailed comparison of CROCHET with existing tools, including MCMICRO^5^, SIMPLI^6^, and TRACERx-PHLEX^7^. The modular structure of CROCHET, organized in three meta-modules, enables streamlined data analyses and supports integration of new methods (**Figure 1**).

**Figure 1.**
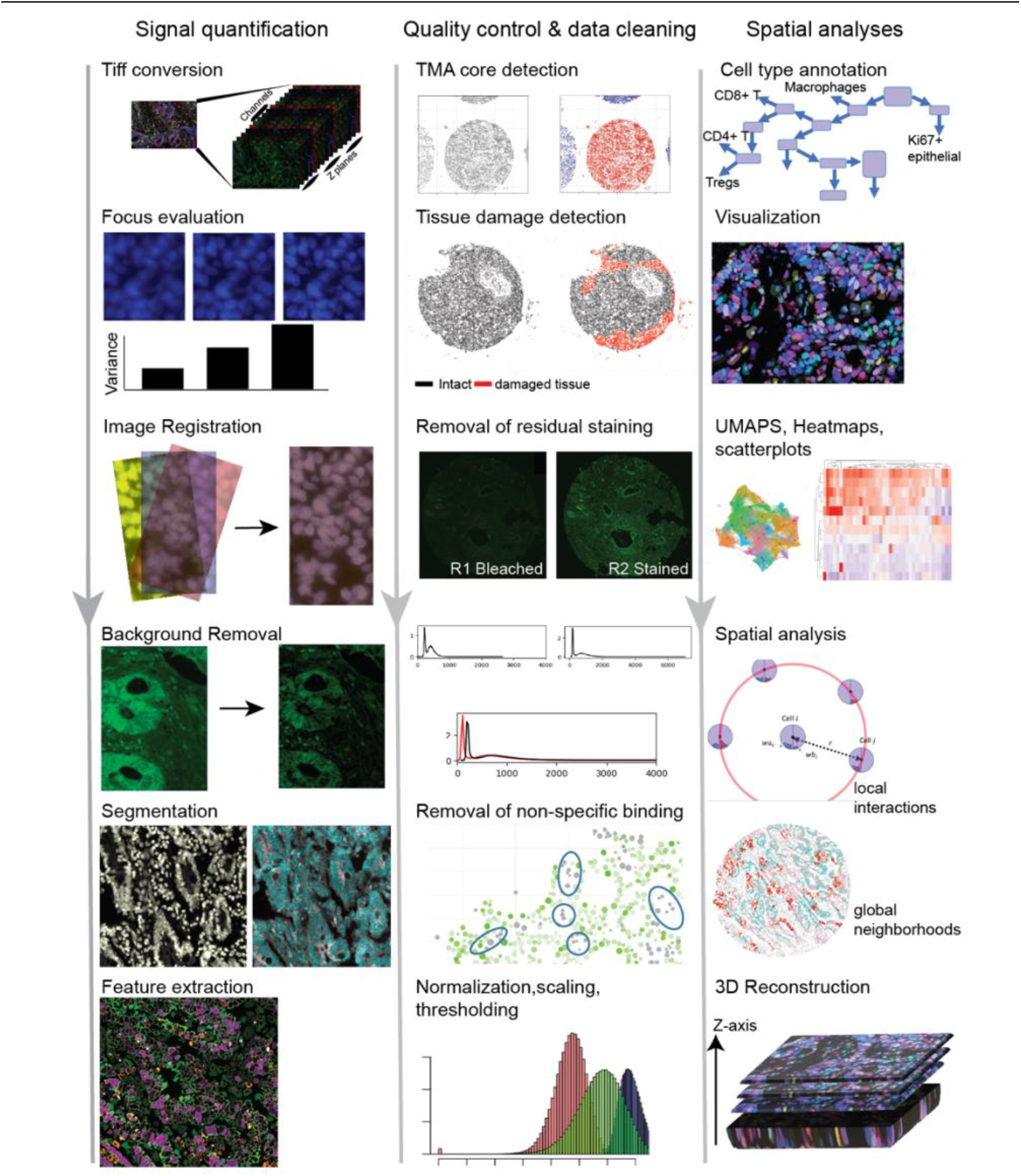
CROCHET spatial omics analysis workflow. The molecular signal quantification module (left) starts with raw data-processing and involves format standardization, focus evaluation, image registration, background estimation, cell segmentation, and feature extraction steps. The quality control and improvement module (middle column) automates the detection of each sample in a tissue microarray, identification of loss/damaged tissue regions, detection and removal of artificial signals, and data normalization. Based on the quality-controlled data, the downstream analysis module (right) annotates cell types using a hierarchical cell phenotyping scheme with well-defined cell type markers, allows interactive visualization of spatial features, single-cell and spatial analyses at both local-interaction and global neighborhood levels and, finally, the construction of 3D tissue maps using data from adjacent tissue slices.

### Meta-module 1

The first meta-module of CROCHET includes image processing, segmentation and feature extraction modules. The image processing module utilizes OME Bio-Formats tools^9^ to support hundreds of image file formats from various instruments and converts them to TIFF, enabling a standardized analysis framework. Next, the focus evaluation step automates the selection of the sharpest Z-sections from each specimen (**Figure 2A, Equation 1**). In the segmentation module, we implemented the CycIFAAP^10^ suite to perform image registration, background removal, and nucleus and cell boundary detection. For registration, images from the same specimen are aligned by superimposing identical nuclei, enabling analyses within a unified coordinate frame. Following background subtraction, images are segmented using the Mask Region-based Convolutional Neural Network (MASK R-CNN) algorithm (**Figure 2B**)^11^. First, nuclear boundaries are segmented based on nuclear staining signal. Next, cellular boundaries are segmented based on the spatial distribution of available cell surface markers around nuclei from a pre-selected set (e.g., E-Cadherin, CD45, CD44). When lacking markers, cell boundaries are estimated by expansion of nucleus boundaries until a neighbor cell is encountered or a maximum cell size is reached (default: 3x nuclear size). Currently, the Cellpose^12^ algorithm is also implemented as an alternative segmentation method. Other segmentation methods can be incorporated, or existing methods can be updated as new modules. Following segmentation, cellular features are extracted from the regions bounded by segmentation masks. These include protein readouts from subcellular regions (cell surface, nucleus, and cytoplasm), spatial features such as cell coordinates, and a range of morphological descriptors. The intensity-based features include protein readouts such as mean, variance, skewness and Kurtosis from subcellular regions (**Figure 2C)**. Spatial features include cell coordinates, bounding box and distance from tissue border. Morphological features include cell size, cell orientation and metrics for symmetry, circularity, directional variance, surface convexity and haralick features (i.e., spatial relations among pixels). For samples processed on tissue microarrays (TMAs), individual tissue sample boundaries are identified using the HDBSCAN algorithm (**Figure 2D**)^13^. The optional normalization steps based on exposure times and light intensity enable comparisons of signals across different imaging cycles (**Equation 2**).

**Figure 2.**
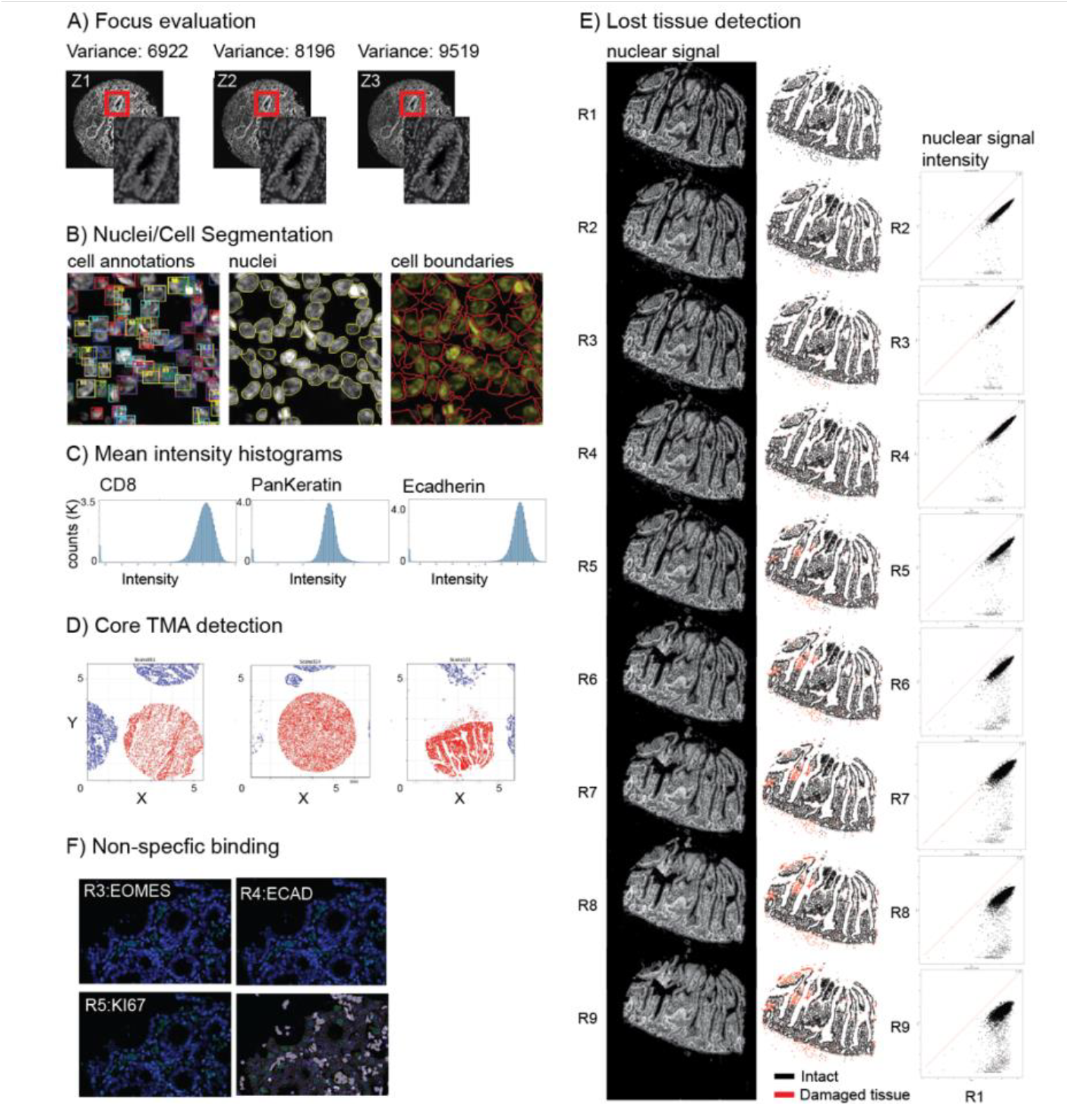
Quantification and quality control of spatial omics data. A) Focus evaluation using normalized variance-based statistical algorithm B) Mask R-CNN single-cell detection and nucleus and cell segmentation algorithm. C) Histograms of protein biomarker mean intensity values within segmented areas. D) TMA core detection using HDBSCAN clustering algorithm. E) Lost tissue detection across experiment cycles via comparison of the nuclear staining intensity values with those obtained in the first cycle. F) Detection of cells with non-specific binding, using protein biomarker expression values across different cycles (See methods for details).

### Meta-module 2

The second meta-module of CROCHET includes quality control and data cleaning modules, leveraging features from the first meta-module or user-provided data. Next, for cyclic imaging methods (e.g., cycIF, COMET, and CODEX), the tissue damage or loss which may be due to cycles of staining and chemical destaining or low sample quality is detected and quantified (**Figure 2E, Equations 3–6**) to evaluate data completeness across imaging cycles. Next, we automate quantification and correction of artificial signals, a major challenge in IF analysis. The form of artificial signal, residual fluorescence from previous staining cycles can contaminate subsequent rounds due to incomplete bleaching. The CROCHET bleach correction module mitigates this problem by estimating carried-over signal intensity using average signal intensity values from bleached images in each channel and retrospectively correcting the data (**Equations 6–8**). The bleach correction step can help avoid harsher and prolonged chemical bleaching and minimizes tissue damage. We have also implemented the nonspecific staining detection/removal module with an adjustable correction function. As nonspecific staining occurs and corresponds to artificially stained structures (e.g., red blood cells, tissue debris) regardless of the protein specific antibodies, such signals recur in identical patterns in multiple consecutive cycles. The nonspecific staining removal module identifies and alleviates the strong recurring signals that are artificially conserved in multiple consecutive staining/imaging cycles within a particular fluorescent channel (**Figure 2F, Equations 10–11**).

### Meta-module 3

The third meta-module leverages the ready-to-analyze (processed, quality-controlled, normalized) data for spatial atlas construction (**Supplementary Table 2**). The modules include the visualization, cell typing, spatial and statistical analyses, spatial neighborhood detection, and 3D tissue reconstruction tools. The visualization module is built on the Napari interactive viewer infrastructure (**Figure 3A**)^14^. The processed data and features are mapped onto tissue images to create an interactive view of cell masks, tissue features, protein/RNA expression levels, and cell states or types. The user interface enables interactive selection of markers, annotation of cell types, and adjustment of threshold parameters to define marker positivity. Cell typing is performed via an interactive Flask-based application that assigns cell identities using a hierarchical gating strategy. Cell types (e.g., epithelial cells, lymphocyte sub-populations, mesenchymal vascular cells) are annotated based on the expression of available cell type markers in the user’s data (**Figure 3B, Supplementary Table 3**). The output is a collection of cells each annotated with a cell-type based on the relevant protein markers.

**Figure 3.**
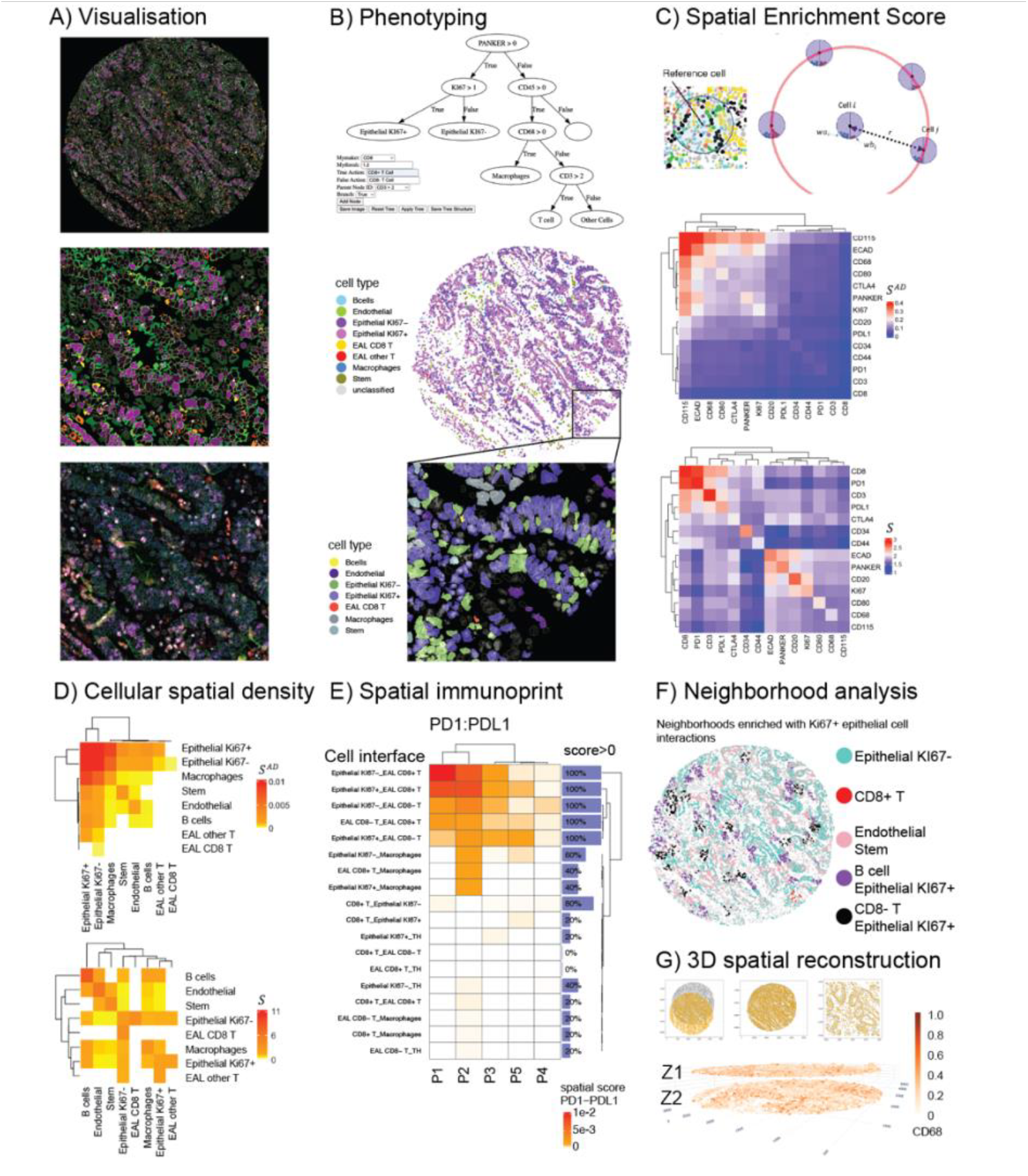
Analysis and visualization of spatial omics data. A) Top: Napari-based visualization of processed data with different colors corresponding to different biomarkers on a gastrointestinal tumor sample. Middle: Mean protein expression values of nuclear proteins are shown within segmented nuclear area and mean protein expression values of non-nuclear proteins are shown within segmented cytoplasmic area. The color intensity represents expression levels. Bottom: Pixel-based images of protein expressions within segmentation masks. B) Top: Interactive phenotyping Flask app to build and apply the hierarchical tree for cell type annotation. A representative tree is shown. Middle: Spatial map of cell types. Bottom: Color-coded visualization of cell types overlying with nuclear and cellular segmentation masks. C) RDF-based spatial enrichment score capturing the expression-weighted enumeration of neighbors in close proximity to a central cell (see methods). The Heatmap for spatial enrichment scores of protein marker pairs without (middle) and with (bottom) global abundance normalization in a whole sample. D) The Heatmap for spatial enrichment scores of cell-cell interactions without (top) and with (bottom) global abundance normalization in a whole sample. E) Immunoprint quantifies PD1-PDL1 spatial interaction on proximal cells across 5 bowel adenocarcinoma patients. F) Mapping of tissue neighborhoods for cell-cell interactions based on Louvain clustering of cell-type spatial enrichment scores at single-cell-level and projection of cluster identities to tissue coordinates. Clusters identify tumor cells with similar neighborhood compositions and interactions. G) Construction of 3D spatial maps through registration and alignment optimization of adjacent tissues.

The spatial analysis module computes a novel spatial enrichment score. CROCHET presents separate modules to quantify spatial scores at single-cell-level and sample-level. CROCHET’s spatial enrichment score quantifies a weighted radial density distribution (RDF) of protein markers or type-annotated cells around an individual cell or within a defined region of interest, resulting in spatial analyses across scales—from an individual cell to entire samples (**Equations 12–20, Figure 3C-D, Supplementary Table 4**). Conventional spatial metrics that are solely based on point density calculations fail to account for varying tissue structures when cellular densities differ substantially. CROCHET overcomes this challenge by normalizing marker or cell-type specific spatial proximity measurements to the local RDF, thereby adjusting for tissue density properties. Normalization of the spatial enrichment score to marker abundance enables spatial analyses that are independent of the overall expression levels, capturing distribution of rare events. As a result, the spatial enrichment score enables direct comparison across different biomarker pairs and effectively captures the spatial co-arrangements, even for sparsely expressed markers or rare cell types — a critical limitation in existing methods. Consequently, the score is also robust to differences in intrinsic architecture of biological tissues. Without normalization, the spatial enrichment score quantifies protein abundance-weighted marker distributions (**Equation 16**). To the best of our knowledge, it is the first spatial metric capable of supporting direct comparisons of cellular organization across heterogeneous tissue types.

To quantify spatial receptor-ligand interactions on proximal cells, we developed a novel framework, termed immunoprint, based on spatial enrichment scores (**Equation 20, Figure 3E**). The immunoprint represents the distribution of receptor-ligand interactions on tissue interfaces across a large cohort of patients similar to the distribution of oncogenic alterations on oncoprints^15^. When used to quantify immune checkpoint co-distributions (e.g., PD1:PDL1) at tumor-immune interfaces, the immunoprint offers a tool for matching individual cancer patients to precision immunotherapy candidates expressed on the surface of specific cell types and tumor-immune interfaces. In addition, CROCHET maps multi-cellular a tissue neighborhoods by carrying multi-variate analysis of the spatial scores at single-cell-level to identify intratissue niches enriched for distinct cell-cell interaction patterns (**Figure 3F, See methods, Supplementary Table 5**).

As an essential step toward constructing cellular interaction maps and neighborhood identification in 3D space, CROCHET incorporates a module that overlays data from adjacent tissue cuts (**Equations 21–22**). By leveraging the coordinates of cells from two adjacent tissue cuts, the pipeline reconstructs 3D tissue continuity across the two data sets using Euclidean transformations. Data from adjacent sets are then spatially mapped using a 3D visualization module (**Figure 3G**).

The CROCHET workflow represents a significant advance toward standardization of spatial analysis of single-cell omics data with particular emphasis on multiplexed cellular imaging independent of experimental method or data modality. It addresses common, persistent challenges in pre-processing, quality control, and analysis of image-based cyclic and fluorophore-based data. Designed as an end-to-end, open source, modular structure, CROCHET enables integration of both existing analysis methods and potential future methods. While broadly applicable across multiple spatially resolved single-cell data types, CROCHET currently does not accommodate data that lack single-cell resolution, such as spot-based low resolution spatial transcriptomics data. Finally, CROCHET spatial enrichment scores provide a uniquely comparable metric across tissue types, suitable for training Artificial Intelligence models for spatial biology.

## Methods

### Data

Tissue microarray (TMA) images were obtained from a cyclic immunofluorescence (CyCIF) experiment consisting of 10 cycles and five channels^8^. For each TMA scene (core), 20 images were acquired in CZI format, corresponding to 10 consecutive staining and bleaching cycles. The images have five fluorescence channels, including Hoechst for nuclear DNA staining (20× magnification, pixel size of 0.33 µm) for the nuclear signal. Stained tissue images were acquired at three focal planes, whereas bleached tissue images were obtained at a single focal plane. Details of the CyCIF experiment, including staining and imaging protocols, tissue type, and antibody list, are provided in Dereli et al^8^. Five images of core samples (scenes) are selected for the purpose of demonstrating the workflow. In cases where mean properties from all TMA samples are needed the cores from Dereli et al^8^ is used.

### Image format standardization and focus evaluation

Multi-channel images are converted to single-channel, single-plane TIFF format using the CROCHET tifconversion module, which utilizes OME Bio-Formats tools^9^. For example, a Carl Zeiss Image (CZI)-formatted image with five channels and three Z-planes is converted into 15 single-plane TIFF images. CROCHET is compatible with 161 OME-supported bio-formats that we convert to TIFF using the tiffconversion module.

To ensure optimal focal selection for downstream analysis. the focus evaluation module assesses image sharpness for images across multiple Z-planes. The focus evaluation module uses normalized variance-based statistical algorithm, selecting the best-focused plane for analysis. The normalized variance, 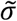, is calculated as the intensity variance over all pixels divided by mean intensity value:

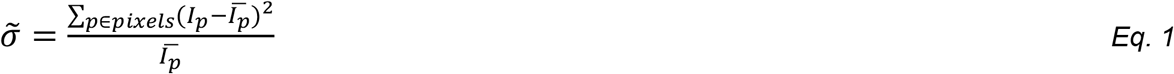

Where *I*_*p*_ is the intensity of pixel p and 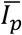 is the mean intensity.

### Image registration background subtraction and segmentation

Images of an individual sample from a multi-cycle experiment are registered by overlaying nuclear signals using OpenCV keypoint detector (SIFT or ORB) and coupled with BFmatcher point matcher. Nuclei segmentation is performed using Mask R-CNN object detection and segmentation framework. We use the trained model provided in cycIFAAP (Cyclic Immunofluorescence Automatic Analysis Pipeline)^10^. The model was trained with 512×512 crops of 5 HER2+ breast cancer TMA cores, as well publicly available immunofluorescence datasets, and achieved 96.9% accuracy and 0.83 dice score on the training dataset. Once the nuclei are segmented, cell segmentation uses nuclear masks and expression of user-selected membrane markers to identify cell boundaries. Once single-cell masks are constructed, mean intensities for each biomarker are calculated in four different sub-cellular zones within cell mask: Nuclei, membrane, cytoplasm, the whole cell. For each single cell, biomarker expressions and spatial features are extracted from each zone, together with spatial coordinates of cell center and its size. Once expressions are corrected and cleaned, users can choose one or more subcellular locations for each biomarker based on the protein’s intrinsic localizations within cell area. CROCHET downstream analysis is independent of a particular image segmentation method. We encourage users to develop and test their segmentation models based on their experimental data. The users can also incorporate their own segmentation results for downstream analysis in the third module. The minimal requirement for CROCHET downstream and spatial analysis are single-cell protein expressions and spatial coordinates.

### Automated tissue detection

In TMA, split scenes (cores) often include parts of neighboring samples. To separate cores (scene) in TMA data and to make sure all single-cell features are matched to the associated sample, CROCHET TMA_detect module uses HDBSCAN clustering algorithm^13^. The location of centroids of segmented single cell nuclei in the first cycle are used to identify the largest cluster of connected tissue. The module identifies to continuous clusters, maps them to individual cores and outputs spatial maps for visual checks. All HDBSCAN python-based arguments are available for the clustering to obtain the best outcomes. Visual inspection and correction are recommended particularly if individual tissue is spatially fragmented and disconnected.

### Intensity and exposure normalization

The cyclic nature of multiplex tissue imaging experiments may require adjusting exposure times and light intensity values at each cycle and for each channel. As a result, to make biomarker expressions comparable, the optional normalization module normalizes protein quantifications by the light intensity and exposure times of their corresponding channel and cycle in the experiment. The module takes the light intensity and exposure time for each cycle and each channel from users and divides the single-cell protein quantifications with their corresponding values:

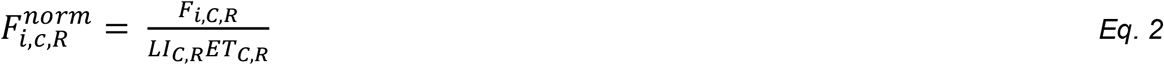

Where *F* _*i,C,R*_ is the biomarker intensity feature from cell *i*, in channel *C*, and round *R*, and *LI*_*C,R*_ and *ET*_*C,R*_ are the light intensity and exposure time of channel *C*, and round *R*.

### Detection of lost cells and tissue damage

Cycles of staining with antibodies and bleaching may result in tissue loss. Here we introduce a module that can detect lost tissue at single-cell-level and remove their corresponding biomarker expression from downstream analysis. To identify lost cells, we quantify nuclear staining (Hoechst) intensity for each single cell as the mean Hoechst staining signal from the nuclear masks. Each cycle of staining and bleaching results in a relative reduction in nuclear staining intensity. We assume that even though nuclear staining intensities decrease in each cycle, they remain correlated to the initial intensity if nuclei remain intact. Figure 2E shows the correlation between different cycles. Once a cell is lost, nuclear staining intensity drops significantly. To identify lost cells in each cycle, we first remove all cells with nuclear staining intensity below a user-defined threshold in that cycle (default 50% of intensity in first cycle). We suggest choosing a non-zero but low value to avoid removing intact cells. For the remaining cells, we calculate their signal ratio across cycles: The ratio between nuclear staining intensity in the corresponding cycle vs. the first cycle. The median and median absolute deviation (MAD) of ratios are calculated:

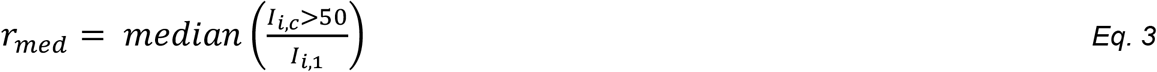

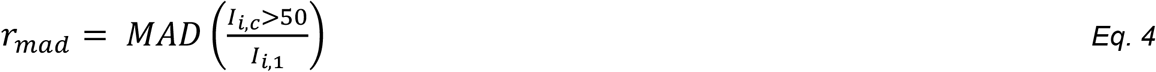

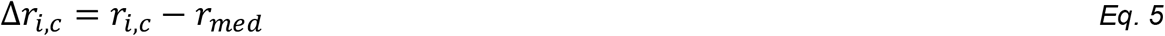

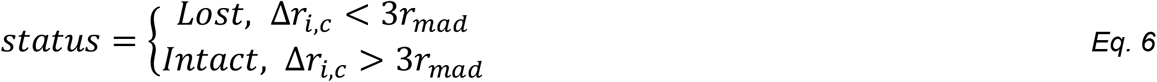

Once the lost cells (Δ*r*_*i,c*_ < 3*r*_*mad*_) are identified, the biomarker expression information for those cells in the corresponding and following cycles are discarded.

### Correction for residual signal from destaining steps

For each stained and destained (bleached) image, we use OTSU method^16^ to separate background noise from foreground tissue-related signal in each channel. The peak locations in intensity histograms of foreground and background are determined and the difference between foreground and background total signal intensity, *I*_*diff*_, (*I*_*diff*_ = ⟨*I*_*foreground*_ ™ *I*_*background*_ ⟩) is calculated. For cases with multiple scenes, as is the case in TMAs, background and foreground differences are averaged over all scenes in TMA. *I*_*diff*_ is then normalized by light intensity (*LI*_*C,R*_) and exposure times (*ET*_*C,R*_) of the corresponding channel (C) and Round (cycle) (R). The ratio *I*_*diff*_ in stained cycle and its following bleached cycle is the estimated fraction of signal that is not properly bleached and is carried over to the next cycle.

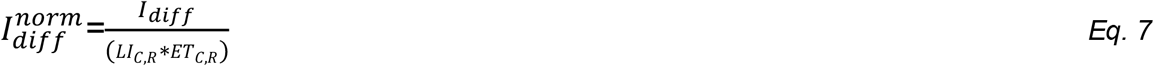

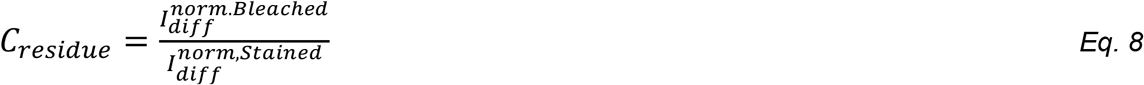

correction coefficient, *C*_*residue*_, is calculated for each cycle and channel. Following segmentation, all single-cell intensity features from segmentation mask is corrected using the coefficient:

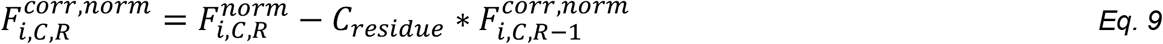

In which 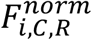 is the biomarker intensity feature from cell *i*, in channel *C*, and round *R* normalized by exposure time and light intensity (Equation 2) from segmented images.

### Filtering non-specific binding (NSB)

One of the main challenges in immunofluorescence-based multiplex imaging is the unwanted binding of primary antibodies to surfaces other than the epitope. Here we provide an automated computational module that identifies likely NSB sites from segmented data and removes them.

First, we have made the observation that nonspecific artificial signals are presented with identical patterns consistently in consecutive cycles with high amplitudes regardless of antibodies, leading to artificial readouts. To decode such artificial signal and resolve, we first identify the brightest signal for each biomarker. Let *I*_*i,c,r,l*_ be the brightness of cell *i* for a given biomarker in channel *c*, cycle *r*, and subcellular localization *l*, and *T* be a biomarker-independent percentile rank (default 85%). A cell is defined as highly bright if:

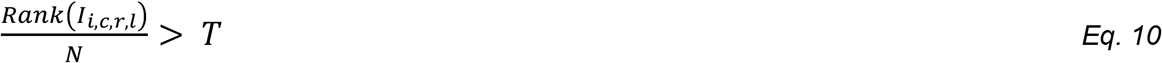

Where *N* is the number of segmented cells in the tissue and *Rank*(*I*_*i,c,r,l*_) is the rank of cell *i* in a list ordered by brightness. Next for each biomarker we check for consistent high ranking, in three previous cycles looking at biomarker intensities in the same channel and same subcellular localization. A staining pattern will be marked as non-specific for a specific biomarker if it satisfies all conditions below:

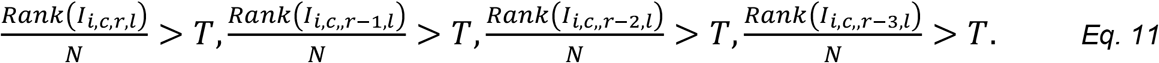

Subsequently, the corresponding biomarker signal for the marked cell is removed from the subcellular localizations. Note non-specific binding sites are marked in subcellular resolution and for a signal to be removed in a specific localization. Therefore, it should rank high in the same exact cellular localizations in previous cycles. As a result, a high intensity of membrane specific biomarker would not interfere with high intensity of a nuclear biomarker. There is a remote possibility that a cell may have extremely high signals for all antibodies used leading to loss of a genuine signal. Yet in most cases, our antibody application design involves antibodies targeting varying cellular localizations in consecutive cycles that minimizes this possibility.

### Cell Type annotations

We present a user-friendly, interactive Flask-based application that enables hierarchical cell classification based on user-defined decision trees. Upon initiation, the application reads expression files. Prior to cell typing, we recommend using CROCHET’s scale function to log normalize and Z scale each biomarker across all scenes. Note that for cases with multiple samples (e.g., cores on a TMA), Z-scaling uses mean and standard deviation of biomarker expression of cells across all samples. This helps applying a consistent threshold value for each marker valid across all samples. Once data is provided, CROCHET displays available biomarker names in a drop-down menu, allowing users to select biomarkers of interest, define threshold values, and assign cell types in a stepwise manner, starting from the root node. The evolving tree structure is visualized at each step (**Figure 3B**) to facilitate intuitive decision-making. Once constructed, the classification tree can be saved and applied to new datasets for automated single-cell annotation. To construct cell types in figure 3B, corresponding biomarker expressions median shifted, log2 normalized, and Z scaled using all TMA scenes (cores) in Dereli et al^8.^

### Radial distribution function (RDF)

The spatial organization of cells are quantified using the RDF formulation. For a given cell *i*, RDF at distance, *r*, is defined as

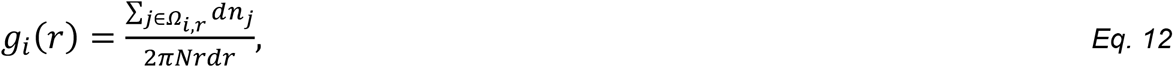

where *N* is the total count of cells in tissue and 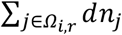 represents the total count of cells within *Ω*_*i,r*_, a shell of thickness *dr* around cell *i* at distance *r*. The CROCHET get_rdf modules computes mean value of *g*_*i*_(*r*) for a user-provided bin-size. While the RDF itself is independent of cell types and expression profiles, we introduce an RDF-based spatial score to build a novel bio-marker dependent metric (see below). The spatial score can be readily extended to quantify the proximity of different cell-types as well as the spatial colocalization of ligand-receptors expressed on different cell types (see immunoprint section).

### Single-cell spatial score for biomarker expression

We first define a weighted radial-distribution-function-based for biomarker expression levels. For each pair of biomarkers *a* and *b*, the weighted RDF, *g*_*ab,i*_(*r*), within a cell neighborhood is given by:

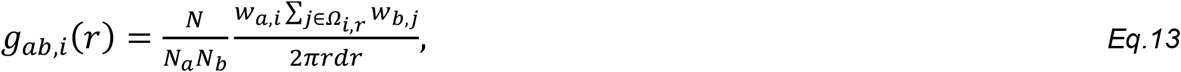

where *W*_*a,i*_ represents weight of marker *a* on cell *i* and is defined as

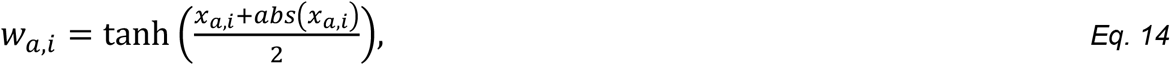

Where *x*_*a,i*_ is the log2-transformed and z-scaled expression of marker *a* on cell *i*. Similarly *W*_*b,j*_ represents weight of marker b on cell *j*, a cell within the shell of thickness, *Ω*_*i,r*_ around cell *i* in distance, r. *N*_*a*_ = Σ_*i*_ *W*_*a,i*_ denotes the total weight of marker *a* in the tissue. Pease note that similar to *g*_*i*_(*r*), integrating *g*_*ab,i*_(*r*) over the entire tissue is constant, making it independent of marker expression levels.

We define the spatial score for cell *i*, biomarkers *a* and *b*, and distance *r, S*_*i,a,b*_(*r*), as the ratio between the integrated weighted RDF for the marker pairs and integrated unweighted RDF for all cells within the radius of r:

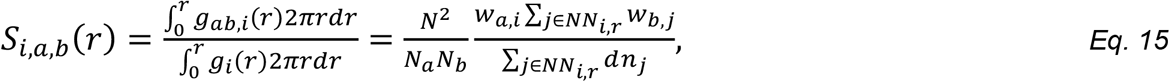

where *NN*_*i,r*_ is the neighborhood of cell *i*, defined as a circular region of radius *r* centered at cell *i, N*_*a*_ and *N*_*b*_ are the total weight of markers for *a* and *b* in the tissue

The *S*_*i,a,b*_(*r*) quantifies the spatial enrichment at the level of individual interacting cells with annotations of cell types or markers. To give users flexibility to quantify both spatial proximity and biomarker abundance in a single measure, CROCHET also provides an abundance-dependent spatial score defined as:

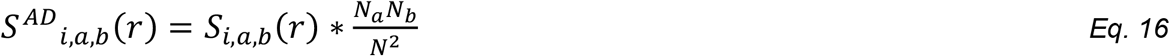

This modified spatial score is useful for users seeking to compare differences between biomarker abundance alongside their interactions across different samples.

### Sample-level spatial scores for biomarker expression

To calculate the sample-level spatial score, *S*_*a,b*_(*r*), we first calculate sum of the weighted RDF for the marker pair over all cells. *S*_*a,b*_(*r*) is defined as the ratio between total weighted RDF to the total unweighted RDF:

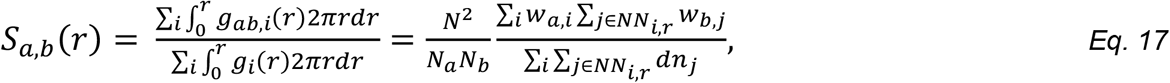

Similarly, the abundance-dependent spatial score at sample-level is:

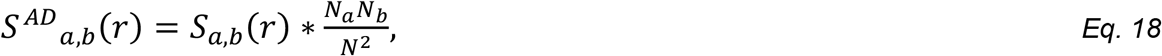

Note that the sample-level *S*_*a,b*_(*r*) does not scale with cell counts and approaches 1 at large distances. This property makes our sample-level spatial score uniquely suitable for direct comparison of spatial scores across different samples and tissues. On the contrary *S*_*i,a,b*_(*r*) scales with cell count and captures local density fluctuations, and for this reason, is best suited in building local niches and neighborhood analysis

### Spatial Score for Cell types

In the same manner as the spatial score for biomarkers, we define a weighted RDF based on cell types. Here we define *W*_*a,i*_ represents cell type of cell *i*:

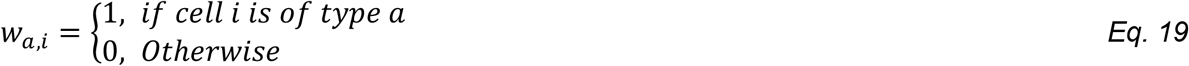

Following the same equations as in biomarker case, the spatial scores to quantify interaction of different cell types at single-cell and sample-levels is calculated using above weights in equations 15–18.

### Immunoprint representation for spatial analysis of ligand receptors interactions

The receptor ligand interactions such as immune checkpoint receptors are modeled using the immunoprint formalism which combines receptor/ligand abundance and spatial distribution on specific cell-cell interfaces. The immunoprints are calculated based on spatial enrichment scores. For immunoprints, weight distances in RDF by both cell types and biomarker expressions are calculated simultaneously. Here, *W*_*a,m,i*_ weight function for cell *i*, cell type *a* and biomarker *m* is given by:

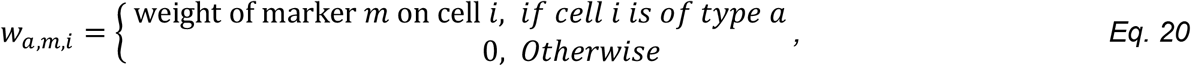

where weights of markers are calculated via equation 14. Immunoprint entry for each sample is quantified by plugging *W*_*a,m,i*_ values in equations 15–18 capturing potential receptor-ligand interactions on cells with specific types within close proximity. The resulting immunoprints represent expression and proximity of ligand/receptors pairs on cells of specific cell types.

### Tissue neighborhood analysis based on spatial interactions

Unsupervised clustering of Z-scaled spatial enrichment scores is performed using the Louvain algorithm via Scanpy^17^ in Python. Depending on the selected spatial score, this approach identifies communities based on the proximities of cell type or biomarker pairs. The neighborhood analysis function inputs cell IDs and their corresponding single-cell spatial scores (for biomarkers, cell types or ligand/receptor pairs) within a tissue sample. First, single-cell spatial scores are log2 normalized and Z scaled. Cells with no valid spatial scores are removed from analysis. A principal component analysis (PCA) is performed using scanpy to compute PCA coordinates and variance decomposition. Ratio of explained variance are plotted for each number of principal components to help users choose the best number of principal components that explains variance ratio. Once users select the number of principal components, nearest neighbor distance matrices are computed using scanpy.pp.neighbors and cells clustered into neighborhoods using the Leiden algorithm (scanpy.tl.leiden). Users can modify the resolution and random seeds for Leiden clustering. The output is a modified input with added Cluster IDs for each single cell as well as full annotated data. Cluster identities are visualized on spatial maps using cell coordinates. Differentially expressed genes for each cluster are identified by default using the Wilcoxon rank-sum test. All optional parameters in Seurat functions remain accessible to users for fine-tuning.

### Visualization

An interactive Napari-based module provides layered visualization to map analysis results on tissue structures. These layers include nuclei and cell masks, processed mean intensity values for each biomarker, background-removed pixel-based images for each biomarker and nuclear staining, cell types, and non-specific binding sites. Users can select specific biomarkers for visualization and interactively toggle layers on or off to create composite images integrating inputs and results. Additionally, the ROI_crop and plot_tissue modules generate spatial maps to visualize selected regions of interest (ROIs). Users can assess tissue damage for quality control. The removal_of lost_cells module outputs intensity correlation scatterplots for each cycle, which can be used for quality control and parameter tuning in tissue damage assessment. Histograms of expression values are provided via the plot_histogram module. For visualization of raw tissue images, we recommend using OMERO Open microscopy environment^18^.

### 3D tissue reconstruction

Images from adjacent tissues are processed and analyzed independently. For each dataset, point densities of cells surrounding each cell is computed at three user-defined neighborhood distances. These point densities are used to construct a feature array for each cell. To map adjacent tissues in 3D continuity, the cosine similarity score for each cell pair in the adjacent and reference tissue is defined as:

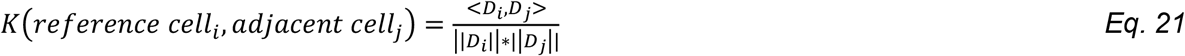

where *D*_*i*_ is the array of point densities (*d*_*i*1_, *d*_*i*2_, *d*_*i*3_) for cell *i* at distances (*r*_1_, *r*_2_, *r*_3_). The highest *K* for each cell in the adjacent tissue corresponds to its most similar cell in the reference tissue. A mapped cell list is constructed from the most similar cells of reference tissue for each cell in the adjacent tissue. Please note that the mapped cell list is built to find possible counterpart cells between the two tissues. As a result, each cell in the mapped list could map to one or more adjacent cells. Mapped list would have the same number of cells as the adjacent tissue, and not all cells in the reference list would be in the mapped list. The mapped list is used to estimate a Euclidean transformation (comprising of rotation and translation) that aligns the coordinates of the adjacent tissue with the coordinates of the mapped list.

This process is iterated, and in each iteration, the Euclidean norm of the difference between the transformation matrix and the identity matrix of the same dimension is computed:

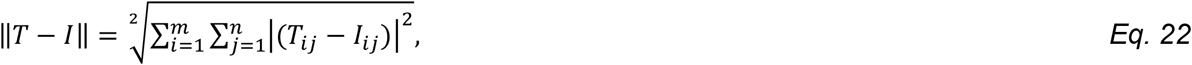

where *T* is the Euclidean transformation matrix and *I* is the identity matrix. As the transformations converge toward the identity matrix, the Euclidean norm decreases, indicating better alignment of adjacent tissue with the reference tissue. Users can modify the neighborhood distances for point density calculations and the number of iterations to tune the outcome for the best alignment.

## Code Availability

CROCHET software and code that is used to create the figures are available.

## Data Availability

Sample SBA images and metadata used to create figures and findings of the study is available at through MD Anderson Data Cloud upon request.

## Acknowledgement

We thank the MDACC BCB software team for support. This work is supported by grants from MDACC Support Grant P30 CA016672, U01 CA253472. This work was supported by the Andrew Sabin Family Foundation, Kavanagh Family Foundation and Kevin T. Doner Memorial Fund (no grant number, philanthropic).

